# TeamTree analysis: a new approach to evaluate scientific production

**DOI:** 10.1101/2020.06.01.128355

**Authors:** Frank W. Pfrieger

## Abstract

Advances in science and technology depend on the work of research teams and the publication of results through peer-reviewed articles representing a growing socio-economic resource. Current methods to mine the scientific literature regarding a field of interest focus on content, but the workforce credited by authorship remains largely unexplored, and appropriate measures of scientific production are debated. Here, a new bibliometric approach named TeamTree analysis is introduced that visualizes the development and composition of the workforce driving a field. A new citation-independent measure that scales with the H index estimates impact based on publication record, genealogical ties and collaborative connections. This author-centered approach complements existing tools to mine the scientific literature and to evaluate research across disciplines.

## Introduction

Progress in science and technology depends on research teams working on specific topics of interest and on the publication of their results in peer-reviewed articles [1]. The rapidly growing body of scientific information [2] reflects past and current states of the art and represents an invaluable socio-economic resource guiding future research activities, policies and investments [3–8]. Its utility relies on the quality and accessibility of bibliographic databases [9, 10] and on refined methods to search and analyse the content of scientific articles [3, 6, 11–16]. Authorship on these articles credits contributions of individual team members with diverse expertise and skills [17–21], but choosing the best method to evaluate research, for example to identify potential experts, recruits and collaborators, remains a challenge [22]. Presently, the impact of individual contributors [23], journals [24], institutions and nations [25] is predominantly estimated based on citation counts of scientific articles (for reviews see [5, 26–28]). In a frequent scenario, a user interested in a specific topic queries a bibliographic database, scrutinizes the resulting list of relevant publications and learns readily about scientific advances. But, it is very difficult for the user to learn about the contributing teams and their impact. To address this recurring issue, I propose a new bibliometric approach, further referred to as TeamTree analysis (TTA). Using author names and publication years of scientific articles related to a field of interest, TTA reveals the development and composition of the workforce with new visuals, named TeamTree graphs (TTGs), and estimates the impact of authors with a new metric named TeamTree product (TTP). TTP takes into account three aspects of scientific production: publication of articles, the generation of offspring and the establishment of collaborations. TTP does not depend on citation counts or journal impact, but scales with the H index [23] and the sum of citations. Here, the principles of TTA are introduced and its main features are illustrated using a generic model and publications from selected fields of science and technology.

## Methodology

The principal steps and key features of TTA are introduced in Fig 1 using generic publications. The TTA-derived parameters are summarized in Table 1. Typically, scientific articles related to a user-defined topic of interest are retrieved from a bibliographic database (Fig 1A; Table 2). From each article, TTA extracts the authors, the year of publication and a database-specific article identifier (Fig 1A). TTA includes author initials to reduce author ambiguity [29]. For some fields, frequent ambiguous author names were removed. TTA categorizes authors according to their byline position and sorts publications by year. Then, it assigns a chronologic author index (AI) and a randomly generated color (C) to each last author (Fig 1B). TTA focuses on authors on the last byline position as they are mostly responsible for the research [19]. In the following, the term “author” refers to “last author” unless indicated otherwise.

**Fig 1.**
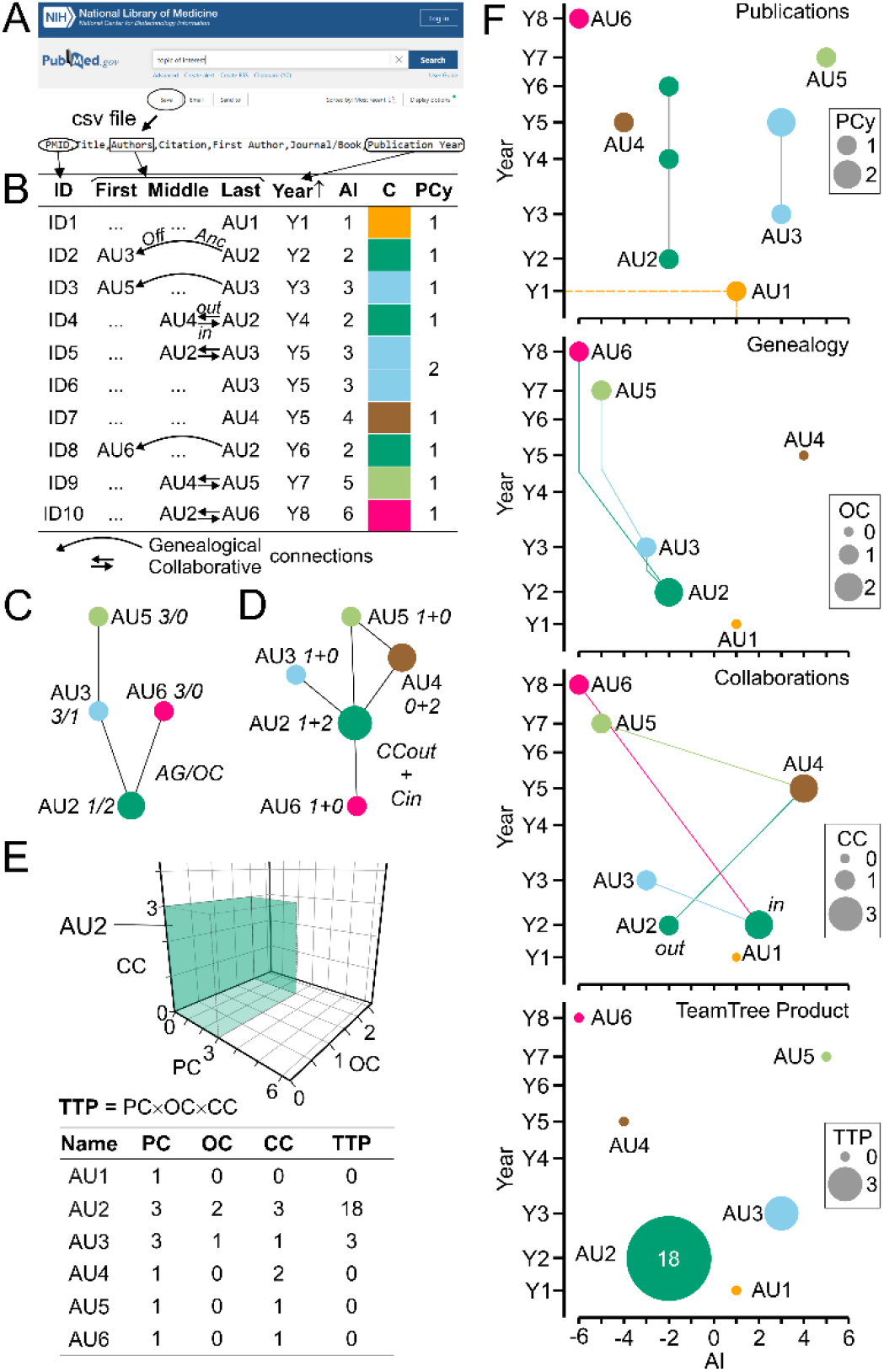
Principal steps and key features of TeamTree analysis. (A) Screenshots of the PubMed website and of a comma-separated values (csv) file illustrating a query in the bibliographic database MEDLINE, the download of scientific articles and the extraction of data required by TTA. (B) Table showing generic articles with identifiers (ID), authors separated by byline position, and years of publication. Only authors mentioned at least once on the last byline position are taken into account and indicated by generic names (AUx). TTA sorts articles by year of publication in ascending order, assigns to each last author a chronologic author index (AI) and a unique color (C) and counts the number of articles per author per year (PCy). Curved arrows indicate genealogical relations between ancestors and offspring on the last and first byline position, respectively. Straight arrows indicate collaborative connections between last authors and co-authors (out) and vice-versa (in). (C) Family tree and (D) collaborative network derived from the generic articles shown in panel B with genealogy- and collaboration-related parameters indicated for each author. AG, author generation; OC, offspring count; CC = CCout + CCin, number of collaborative connections. (E) Three-dimensional plot of key metrics (PC, publication count as last author) for a selected author (AU2) shown in panel B. The volume occupied by the author within the parameter space is indicated by the author-specific color and represented numerically by the TeamTree product (TTP). The table summarizes the TTA-derived parameters of generic authors. (F) TeamTree graphs (TTGs) of the generic authors shown in panel B indicating from top to bottom their publication record, genealogic and collaborative connections and TTP values. For publications and TTP values, signs of AI alternate between odd and even values. For genealogic relations, signs of family members are determined by the first generation author. To indicate collaborative connections, AI of last authors and co-authors are negative and positive, respectively. Symbol sizes represent indicated parameters.

**Table 1.**
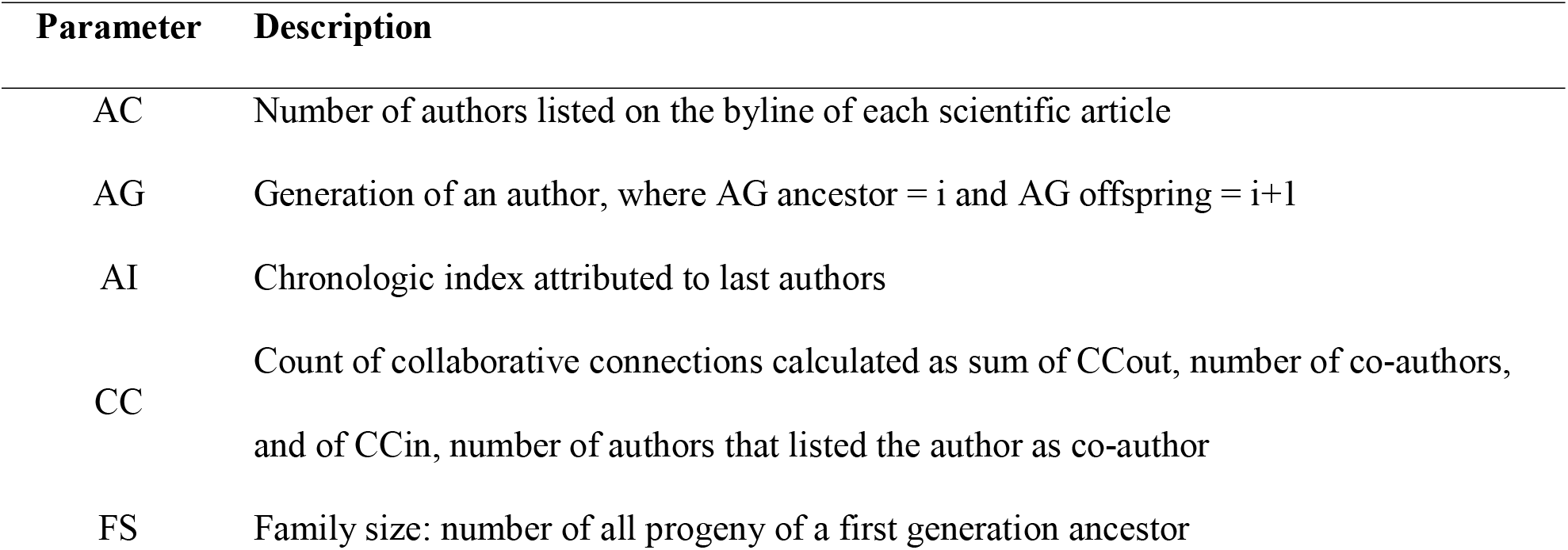
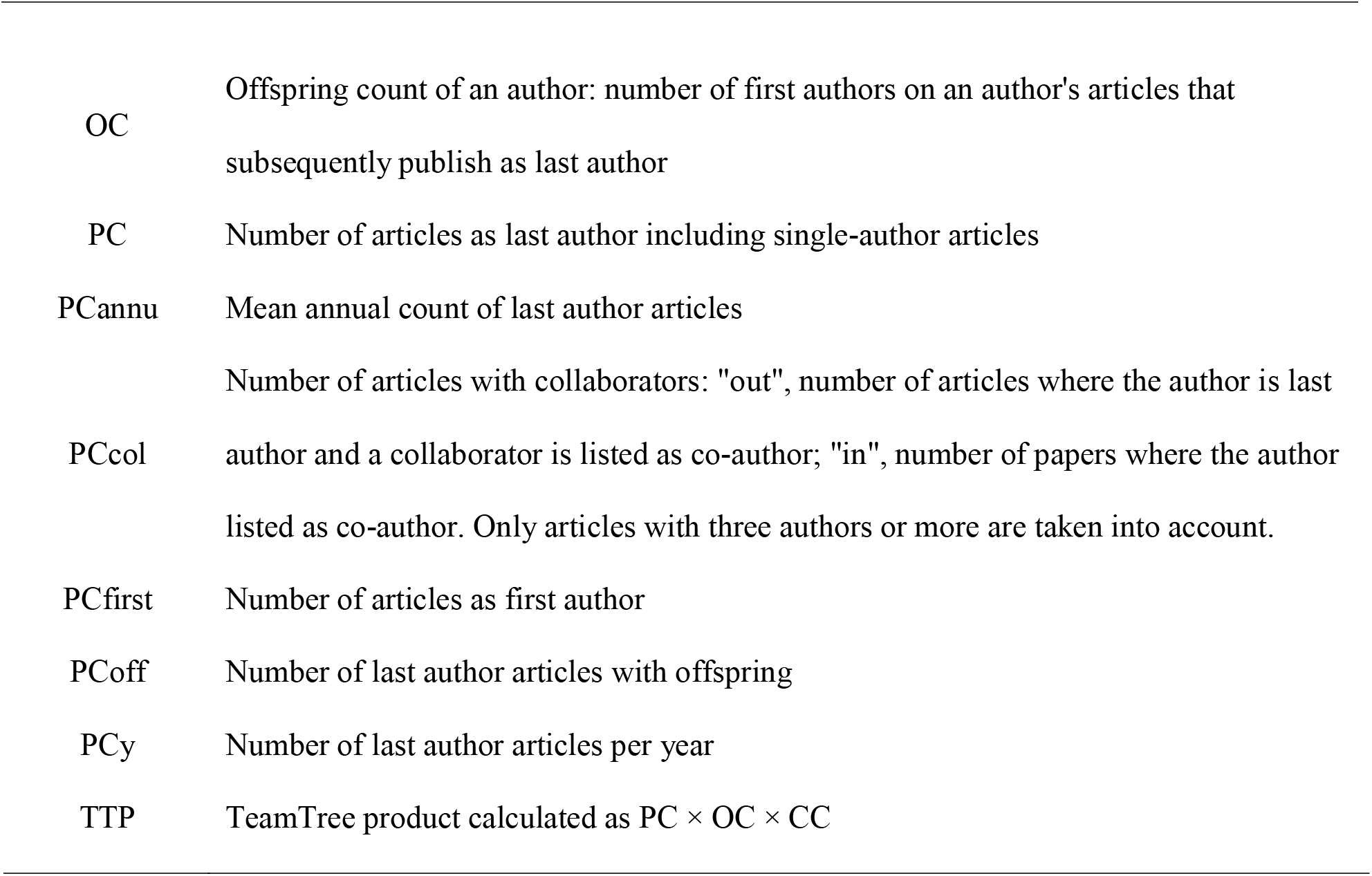
TTA-derived parameters.

| Parameter | Description |
| --- | --- |
| AC | Number of authors listed on the byline of each scientific article |
| AG | Generation of an author, where AG ancestor = i and AG offspring = i+1 |
| AI | Chronologic index attributed to last authors |
| CC | Count of collaborative connections calculated as sum of CCout, number of co-authors, and of CCin, number of authors that listed the author as co-author |
| FS | Family size: number of all progeny of a first generation ancestor |
| OC | Offspring count of an author: number of first authors on an author's articles that subsequently publish as last author |
| PC | Number of articles as last author including single-author articles |
| PCannu | Mean annual count of last author articles |
| PCcol | Number of articles with collaborators: "out", number of articles where the author is last author and a collaborator is listed as co-author; "in", number of papers where the author listed as co-author. Only articles with three authors or more are taken into account. |
| PCfirst | Number of articles as first author |
| PCoff | Number of last author articles with offspring |
| PCy | Number of last author articles per year |
| TTP | TeamTree product calculated as $PC \times OC \times CC$ |

**Table 2.**
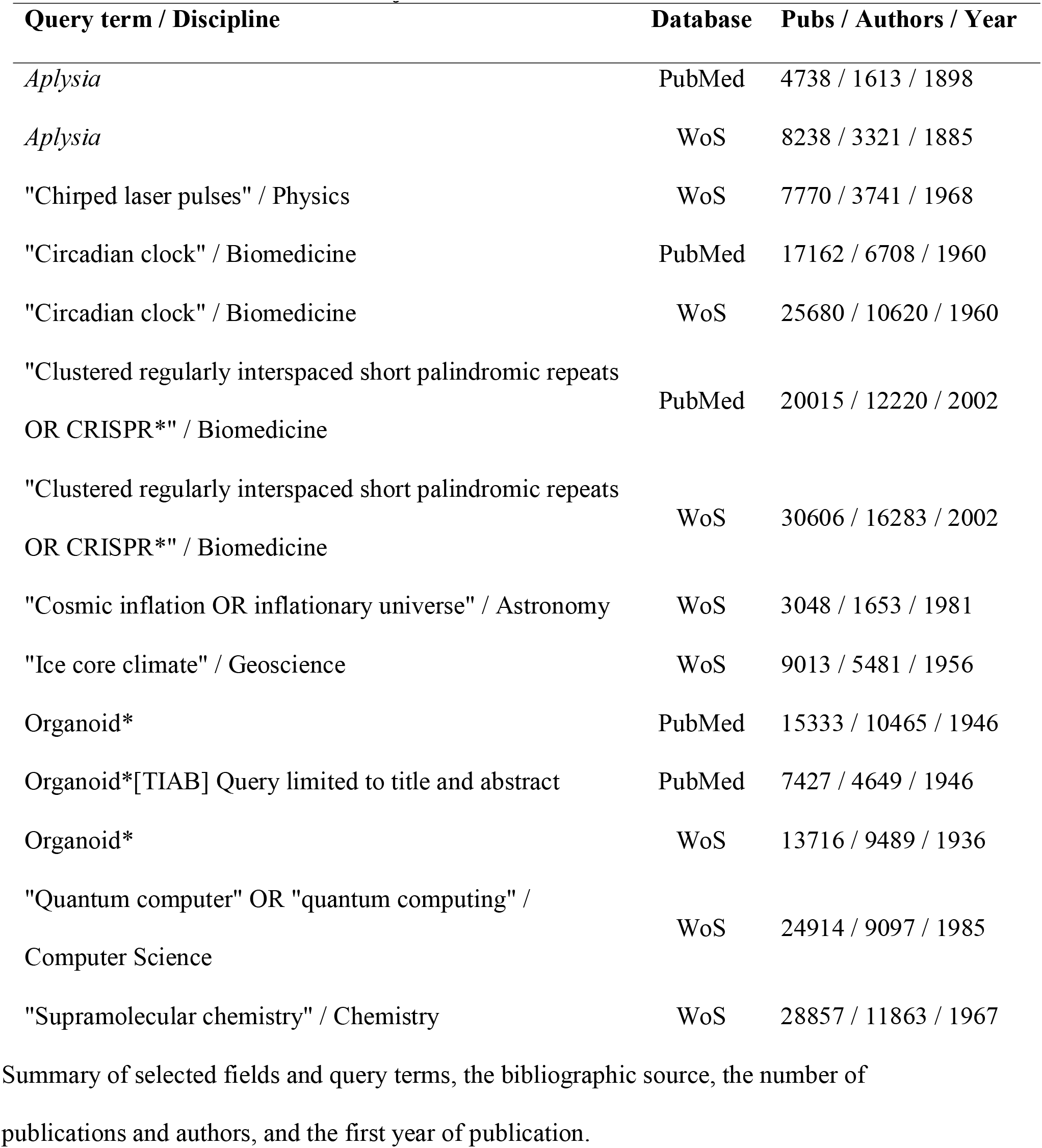
Selected research fields subjected to TTA.

| Query term / Discipline | Database | Pubs / Authors / Year |
| --- | --- | --- |
| *Aplysia* | PubMed | 4738 / 1613 / 1898 |
| *Aplysia* | WoS | 8238 / 3321 / 1885 |
| "Chirped laser pulses" / Physics | WoS | 7770 / 3741 / 1968 |
| "Circadian clock" / Biomedicine | PubMed | 17162 / 6708 / 1960 |
| "Circadian clock" / Biomedicine | WoS | 25680 / 10620 / 1960 |
| "Clustered regularly interspaced short palindromic repeats<br>OR CRISPR*" / Biomedicine | PubMed | 20015 / 12220 / 2002 |
| "Clustered regularly interspaced short palindromic repeats<br>OR CRISPR*" / Biomedicine | WoS | 30606 / 16283 / 2002 |
| "Cosmic inflation OR inflationary universe" / Astronomy | WoS | 3048 / 1653 / 1981 |
| "Ice core climate" / Geoscience | WoS | 9013 / 5481 / 1956 |
| Organoid* | PubMed | 15333 / 10465 / 1946 |
| Organoid*[TIAB] Query limited to title and abstract | PubMed | 7427 / 4649 / 1946 |
| Organoid* | WoS | 13716 / 9489 / 1936 |
| "Quantum computer" OR "quantum computing" /<br>Computer Science | WoS | 24914 / 9097 / 1985 |
| "Supramolecular chemistry" / Chemistry | WoS | 28857 / 11863 / 1967 |

Summary of selected fields and query terms, the bibliographic source, the number of publications and authors, and the first year of publication.

TTA explores three aspects of scientific production: the publication record of authors, their genealogical relations and their collaborations. Several parameters are calculated to assess performance in each category (Table 1). To summarize the publication record of each author, TTA calculates the total numbers of articles listing the author on the first (PCfirst) and last byline position (PC), the number of publications (as last author) in each year (PCy; Fig 1B; Table 1), the publication period in years and the average annual publication count (PCannu; Table 1). Single author articles are counted as last author publications. Genealogical relations between authors are derived from offspring – ancestor pairs, where offspring and ancestor are listed on the first and last byline position of an article (Fig 1B, C). Three conditions apply: First, each offspring is assigned to a single ancestor with the earliest common article defining a genealogical relation. Second, this common article has to be published before the earliest (last author) publication of the offspring. Third, the AI value of the ancestor must be smaller than the one of the offspring. TTA assigns a generation index (AG) to ancestors (AG = i) and offspring (AG = i+1; Fig 1C; Table 1) and calculates for each ancestor the number of offspring (OC; Fig 1C) and the number of articles published with offspring (PCoff; Table 1). Families are defined as progeny of a first generation ancestor (AG = 1) encompassing all offspring (AG > 1). TTA derives collaborations based on co-authorship [30] (Fig 1B, D). For out- and in-degree connections, an author lists other authors as co-authors and an author is listed as co-author, respectively (Fig 1B). TTA calculates the numbers of these connections (CCin, CCout; Fig 1C), their sum (CC = CCin + CCout) and the number of corresponding publications per author (PCcol; Table 1). The TTA-derived metrics – PC, OC and CC – define a three-dimensional space, in which each author occupies a distinct volume reflecting publications, offspring and collaborative connections (Fig 1E). The product of these parameters, further referred to as TeamTree product (TTP), defines a new metric to estimate author contributions to a research field (Fig 1E; Table 1).

The workforce contributing to the field is visualized by TTGs. TTGs are scatterplots where each author is represented by a symbol with the AI value and the earliest year of publication plotted on the x and y axis, respectively. The symbols are displayed with author-specific colors (Fig 1F). TTGs provide a framework to illustrate an author’s contributions to each category analysed by TTA. To show the publication records, symbols connected by lines represent the years of publication with symbol sizes indicating the number of articles per year. To achieve an accessible presentation of the publication data, the sign of AI values alternates between odd (positive) and even (negative) numbers rendering a symmetric tree-like design (Fig 1F). Genealogical relations between authors are indicated by lines connecting ancestors and offspring. To represent this aspect with TTGs, the sign of the AI representing the first generation ancestor determines the AI sign of all family members (Fig 1F). To visualize collaborations in the field, lines connect last authors and co-authors with AI signs adjusted to negative and positive values, and symbol sizes indicating CCout and CCin values, respectively (Fig 1F). To represent the overall contribution of an author to the field, TTGs show authors with alternating AI signs and symbol areas representing TTP values (Fig 1F).

TTA is implemented with custom-written routines based on the open source software R [31] and selected R packages for data handling (data.table [32]), statistical and network analyses (igraph [33]; dunn.test [34]) and data visualization (eulerr [35]; ggfortify [36]; ggplot2 [37]; ggrepel [38]; igraph [33]; plot3D [39]). The R script is freely available upon request to the author and at https://github.com/fw-pfrieger/TeamTree. It can be used to analyse publications in a user-defined field of interest. Bibliographic records were obtained from MEDLINE using PubMed (https://pubmed.ncbi.nlm.nih.gov/) and from Web of Science (WoS) (https://apps.webofknowledge.com/; accessed via institutional subscription). To compare citation-independent TTP values with citation-based metrics, the Hirsch indices and the total number of citations were calculated from bibliographic records (WoS).

## Results

To expose the utility of TTA, the new approach was applied to scientific articles from selected fields of research in science and technology (Table 2).

### Visualizing the workforce driving research fields

A new type of visual named TTG reveals the ensemble of authors contributing to a topic of interest (Fig 1). To exemplify this, TTA was applied to three fields of biomedical research each of which showing distinct history, size and dynamics (Fig 2). Corresponding publications were obtained from PubMed/MEDLINE (Table 2). Research on *Aplysia*, a genus of sea slugs, started at the end of the 19th century. Since then, the field expanded slowly but steadily reaching less than 2000 authors total [40] (Fig 2A). The discovery of “clustered regularly interspaced short palindromic repeats” (CRISPR) and the subsequent development of CRISPR-derived genetic tools established a new field, whose workforce is expanding exponentially reaching more than 10,000 authors within a decade [41] (Fig 2B). The field related to “organoids” shows a peculiar development. The workforce expanded transiently during the 1970ies and much of the 80ies (Fig 2C), but this phase was probably due to changing definitions of the term and its assignment to publication records [42]. It is absent when only publications bearing the term in the title or abstract are taken into account (Fig 2C; Table 2). The exponential growth of the workforce within the last decade (Fig 2C) was driven by important breakthroughs suggesting organoids as models of human organs [43, 44].

**Fig 2.**
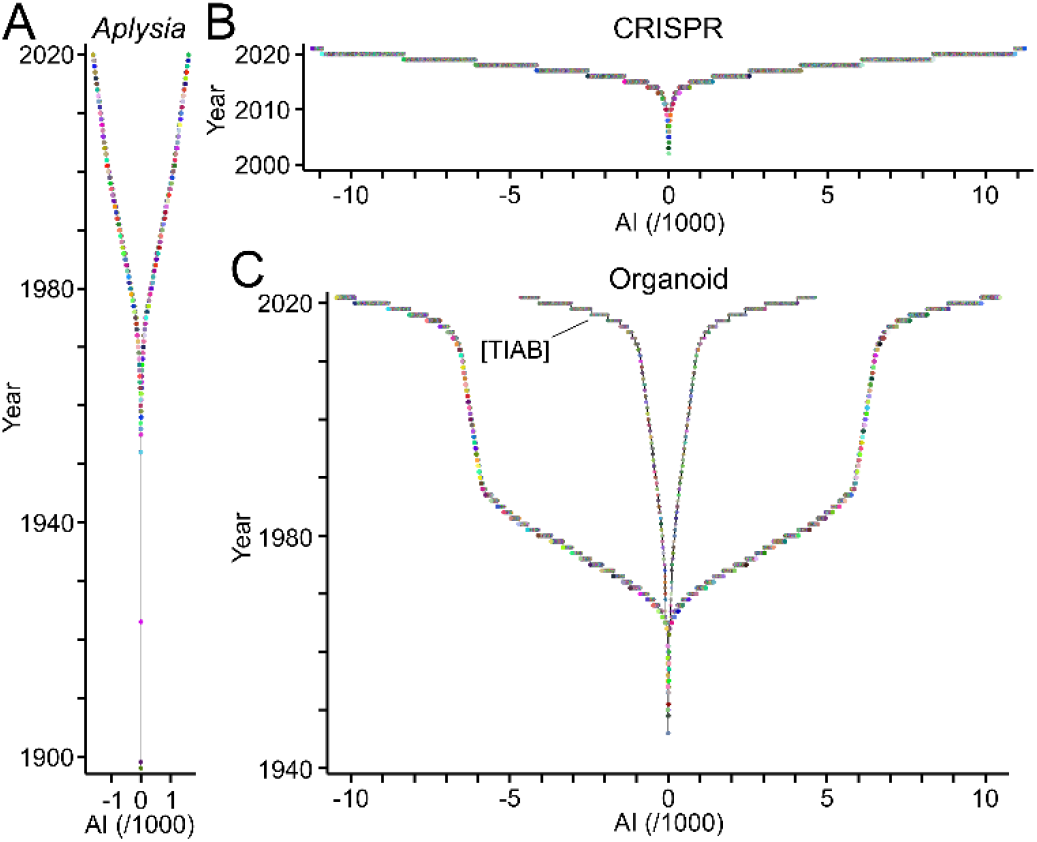
TeamTree graphs showing the development of selected fields of biomedicine. TTGs reveal the distinct duration, growth and size of the workforce publishing scientific articles related to *Aplysia* (A), CRISPR (B) and organoids (C). Circles represent authors contributing to each field with the year of their first publication as last author plotted against their AI values. Signs of AI values alternate for better accessibility. Note the distinct development of the “organoid” field in panel C when only publications were analysed, where the term “organoid*” is only mentioned in the title or abstract as indicated by the field specifier [TIAB].

### Display and quantitative analysis of publication record, genealogy and collaborations

TTA evaluates the publication record of authors, the generation of offspring and the establishment of collaborations. To illustrate this point, TTA was applied to publications related to “circadian clock” (Clock) [45], a well-established field of biomedical research (source: PubMed/MEDLINE; Table 2). Fig 3 shows the publication records of authors in the Clock field using TTGs as framework. Individual authors published as many as 120 articles (PC), but 70% of the workforce contributed single articles (Fig 3B). This percentage was similarly high (68%), when authors entering during the last two years were excluded. The Clock field expanded rapidly within the last decades as indicated by linearly growing annual counts of newly entering authors and of published articles per year, respectively (Fig 3C). Ranking authors by PC values identified the top contributors of articles to the Clock field (Fig 3D).

**Fig 3.**
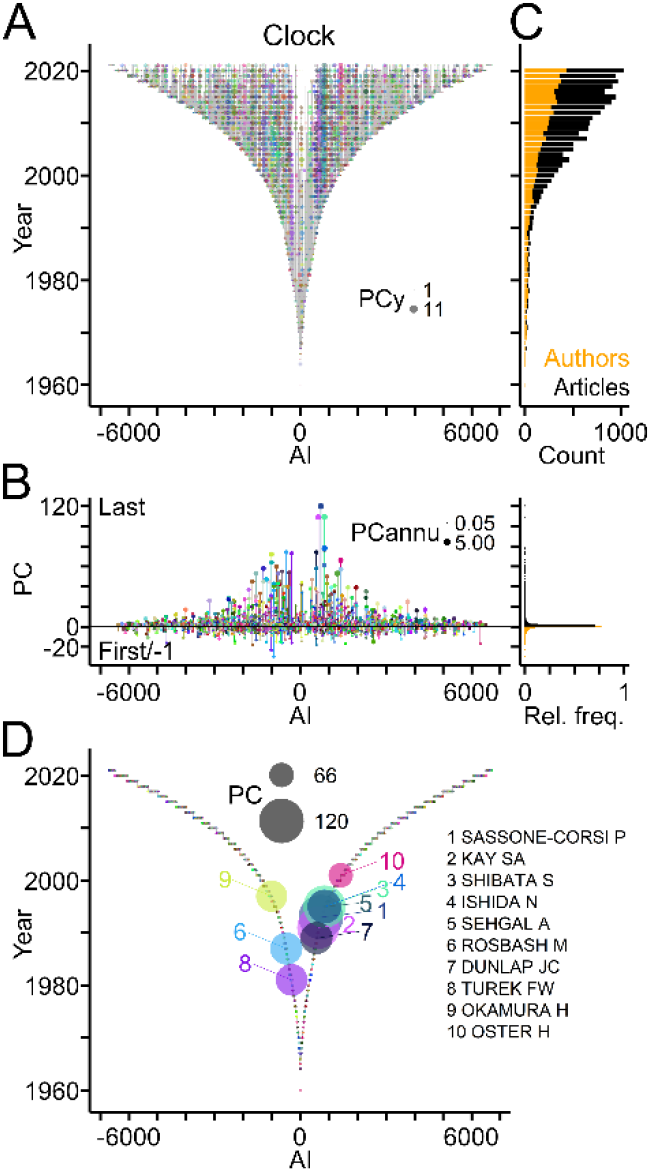
Publication records in the Clock field. (A) TTG showing the publication records of authors working in the Clock field. Circles connected by vertical grey lines represent for each author, the years of publications as last author plotted against the AI. Circle area indicates number of publications per author per year (PCy). (B) Left, publication counts per author indicating last and first author articles by positive and negative values, respectively. Circle area indicates the average number of publications per year (PCannu). Right, relative frequency distributions of PC values shown on the left. (C) Number of authors entering the field per year (orange) and of articles (black) published per year. (D) TTG showing authors with top ten PC values indicated by circle area.

Fig 4 depicts genealogical relations in the Clock field based on last author - first author pairs of articles, and presents a quantitative assessment (Table 1). A quarter of authors published previously as first authors thus qualifying as offspring (Fig 3B) and 10% of the authors qualified as ancestors (Fig 4B). Ancestors generated up to 24 offspring and published up to 75 articles with their offspring (Fig 4B). Overall, the Clock field comprised 506 families with up to 40 members spanning maximally 6 generations (Fig 4B). For the last two decades offspring authors and publications with offspring represented a small, but constant fraction of the workforce entering the field each year and of the annual scientific production (Fig 4C). Ranking by OC values revealed the most prolific authors and their families in the Clock field (Fig 4D).

**Fig 4.**
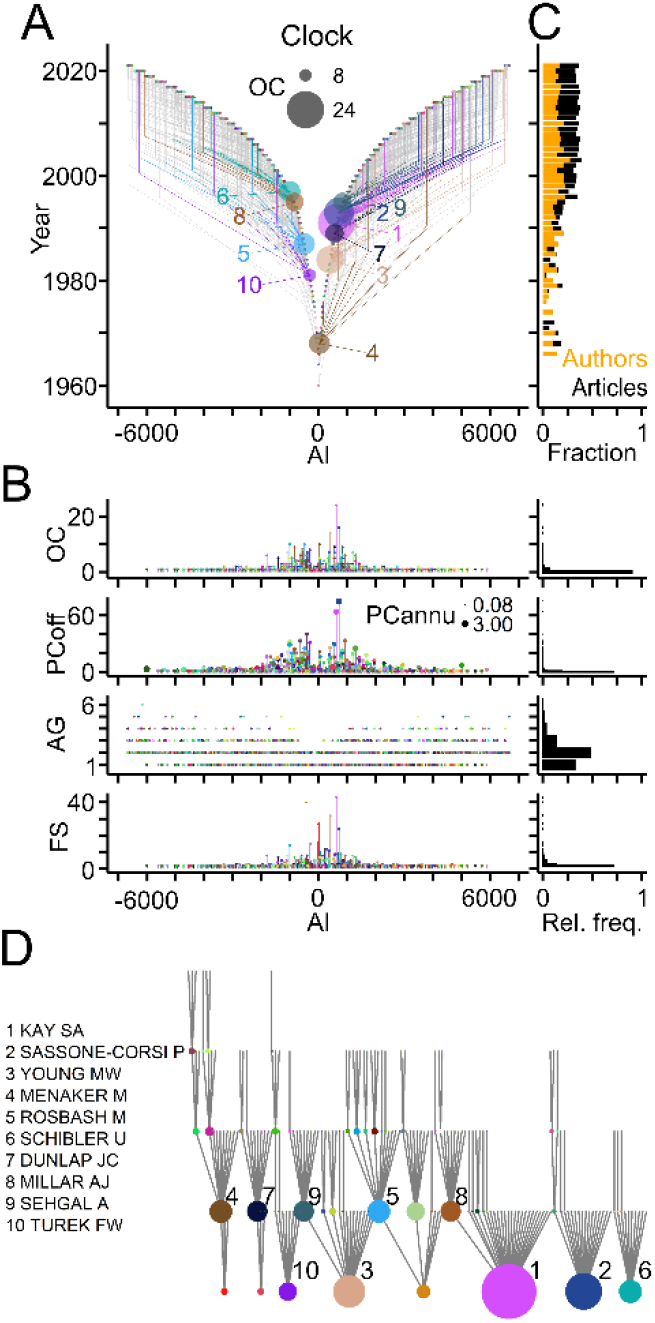
Genealogical relations in the Clock field. (A) TTG showing genealogic relations derived from publications. Circles and grey lines indicate ancestor-to-offspring connections. Connections of authors with the ten largest offspring count (OC) values are shown in color (names indicated in panel D). Circle area indicates OC. AI signs of offspring and of ancestors were adjusted to the first generation ancestor. (B) Left, from top to bottom, OC values, number of articles with offspring (PCoff), author generation (AG) and family size (FS). Circle area indicates PCannu. Right, relative frequency distributions of parameters shown on the left. (C) Fraction of offspring authors (orange) entering the field and of publications with offspring (black) compared to total numbers per year. (D) Names and family connections of authors with top ten OC values indicated by circle area.

Fig 5 shows collaborative connections in the Clock field based on co-authorship and quantitative data using collaboration-specific parameters (Table 1). In total, half of the authors in the Clock field established a variable number of out- and in-degree collaborations with up to 90 authors and published up to 104 collaborative papers as last and co-author, respectively (Fig 5B). During the last two decades, collaborators represented half of the new authors entering per year and their contribution remained fairly constant (Fig 5C). The number of authors per article increased steadily (Fig 5A). Ranking authors based on collaboration counts revealed strongly connected teams in the field and their networks (Fig 5D).

**Fig 5.**
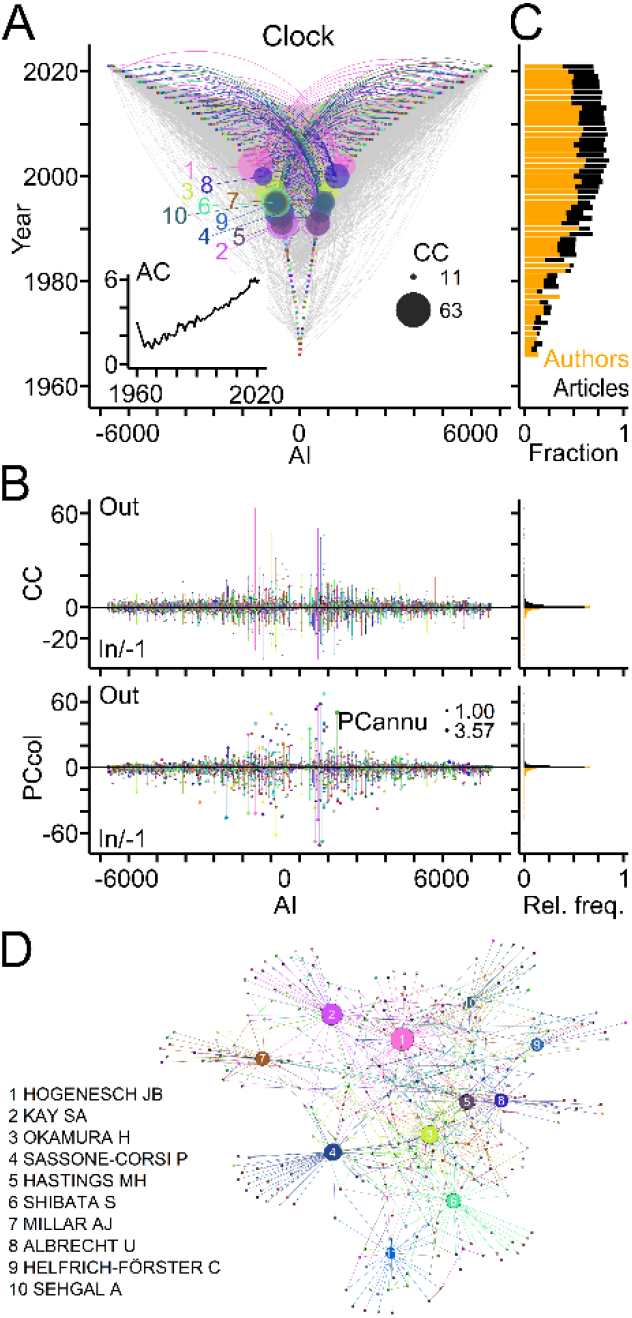
Collaborative connections in the Clock field. (A) TTG showing collaborations between last authors (out; negative AI) and co-authors (in; positive AI) derived from co-authorship on scientific articles. Connections of authors with ten highest connection count (CC) values (in+out) are shown in color. Circle areas indicate CCout and CCin values of these authors. Inset shows the mean author count (AC) per article published each year. (B) Left, counts of collaborators and of collaborative articles per author. Circle area indicates PCannu. Right, relative frequency distributions of parameters shown on the left. (C) Fractions of new collaborating authors (orange) and of collaborative publications (black) compared to total numbers per year. (D) Names of authors with top ten CC values and their networks. Circle area indicates CC values normalized to the maximum.

### Workforce dynamics and field development

TTA was used to explore how the workforce of the Clock field developed over time. Plotting the number of authors entering and exiting the field based on the first and last year of their publications, respectively, indicated strong growth of the workforce. The accuracy of exit counts decreases for the last years (Fig 6). The publication periods or life-spans of authors reached nearly five decades, but the large majority published only during one year and in most cases a single article (Fig 3C; Fig 6A-C). Separating “Newcomers” entering the field per year from “Established” authors revealed that the established workforce consisted mostly of authors with genealogical and collaborative ties, whereas most newcomers had collaborative connections or no ties and contributed single articles (Fig 6D).

**Fig 6.**
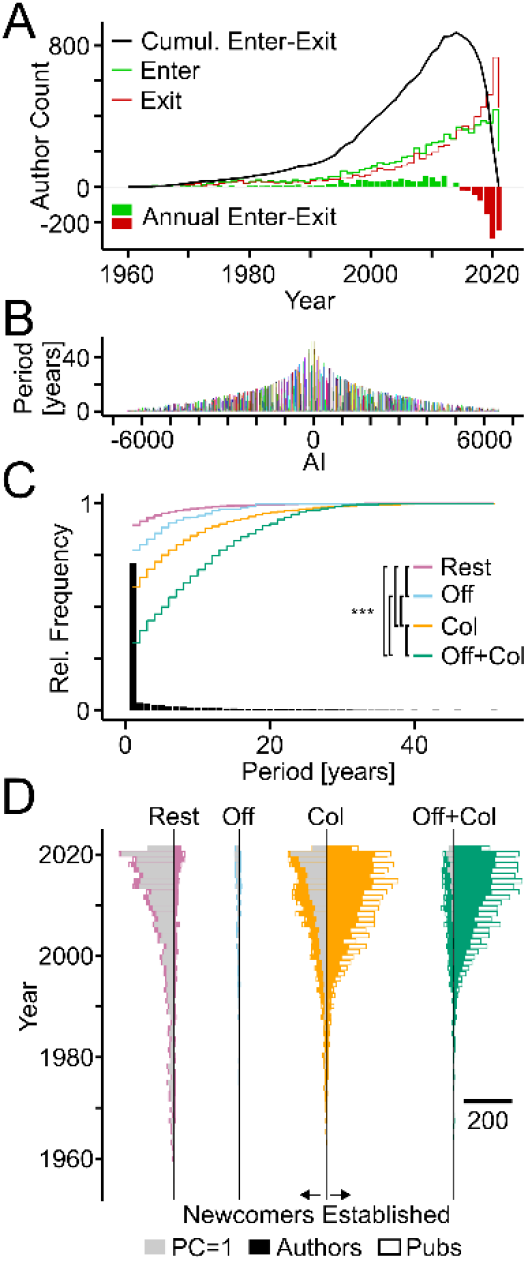
Workforce dynamics in the Clock field. (A) Annual counts of authors entering (green bars) and leaving the field (red bars) with lines showing cumulative sums. (B) Publication periods of individual authors in years. (C) Bars and lines showing the relative frequencies of all publication periods and the cumulative relative frequencies of publication periods of authors from indicated categories, respectively. Col, authors with collaborative but no genealogical connections; Off, genealogical but no collaborative connections; Off+Col, both types of connections; Rest, without connections. Statistically significant differences among groups are indicated (Kruskal-Wallis tests chi-squared = 265.12, df = 3, p < 0.0001. Asterisks indicate level of significance: ***, p < 0.001; post-hoc Dunn test, Benjamini-Hochberg adjusted; sample size = 256; adjusted to smallest sample size by random selection). (D) Horizontal bars indicate number of authors (filled) and of publications (white) per year of newcomers (left) and established teams (right) from the indicated categories. Grey bars indicate authors with single publications. Scale bar indicates number of authors and publications.

### Evaluation of scientific production based on publications, offspring and collaborations

A key goal of bibliometric analyses is to gauge author impact on a field of research. The new metric TTP takes into account an author’s publication record (PC), offspring generation (OC) and collaborations (CC) (Table 1). The concept was introduced with generic publications (Fig 1). Its validity was tested first using publications related to the Clock field (Fig 7). Intersection of the top 100 authors ranked by three key parameters showed that a core of 43 authors figured among the top in all three categories (Fig 7A). Three-dimensional scatterplots of the parameters revealed that authors occupy distinct volumes (Fig 7B) indicating that TTP, calculated as product PC × OC × CC, allows for a more differentiated author ranking than each parameter alone. Fig 7C shows authors with top ten TTP values in the Clock field. To validate its utility, TTP was compared with frequently used citation-based benchmarks of author performance. Scatterplots and statistical analyses revealed that TTP values of individual authors working in the Clock field correlated with the total numbers of citing articles (ρ = 0.828; p < 0.001) and with their H indices (ρ = 0.924; p < 0.001; n = 731; Spearman’s rank correlation; Fig 7D).

**Fig 7.**
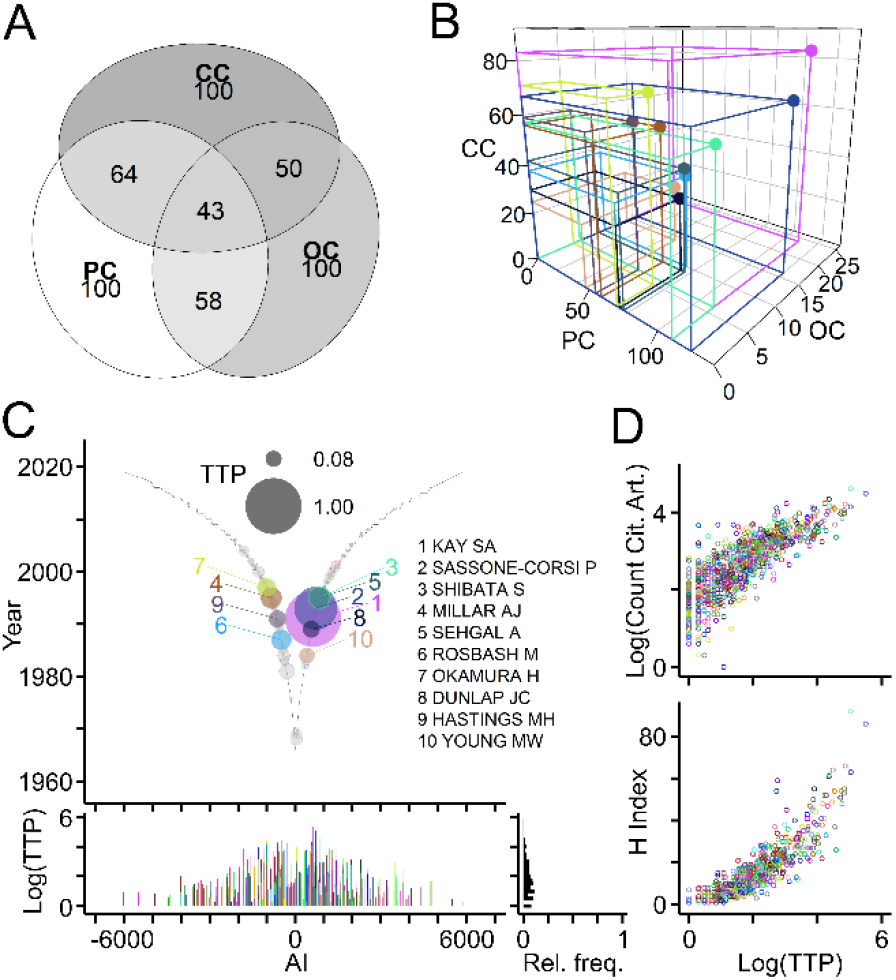
Introduction of TeamTree product as new measure of scientific production. (A) Numbers of intersecting authors in the Clock field ranking among top 100 for each parameter (PC, OC, CC). (B) Scatterplot of indicated parameters for authors with top ten TeamTree product (TTP) values calculated as the volume occupied by each author (PC × OC × CC). (C) Top, graph showing the TTP of authors in the Clock field with colored circles and names indicating authors with ten higest values. Grey circles with colored border indicate authors with TTP values above zero. Circle size indicates log10(TTP) normalized to maximum. Bottom, log10(TTP) values and their relative frequency distribution. (D) Scatterplots, where circles represent individual authors (indicated by color) with their total number of citing articles (top; log10 values) and their H indices (bottom) plotted against their TTP (log10 values).

To further validate TTP as citation-independent measure of productivity, TTA was applied to publications from the fields of biomedical research shown in Fig 2 and to selected fields of science and technology (Table 2). As shown in Fig 8, the TTP values of authors correlated significantly with their H indices and citation counts across fields and disciplines (Fig 8A), and ranking authors by TTP values identified key players in each field (Fig 8B).

**Fig 8.**
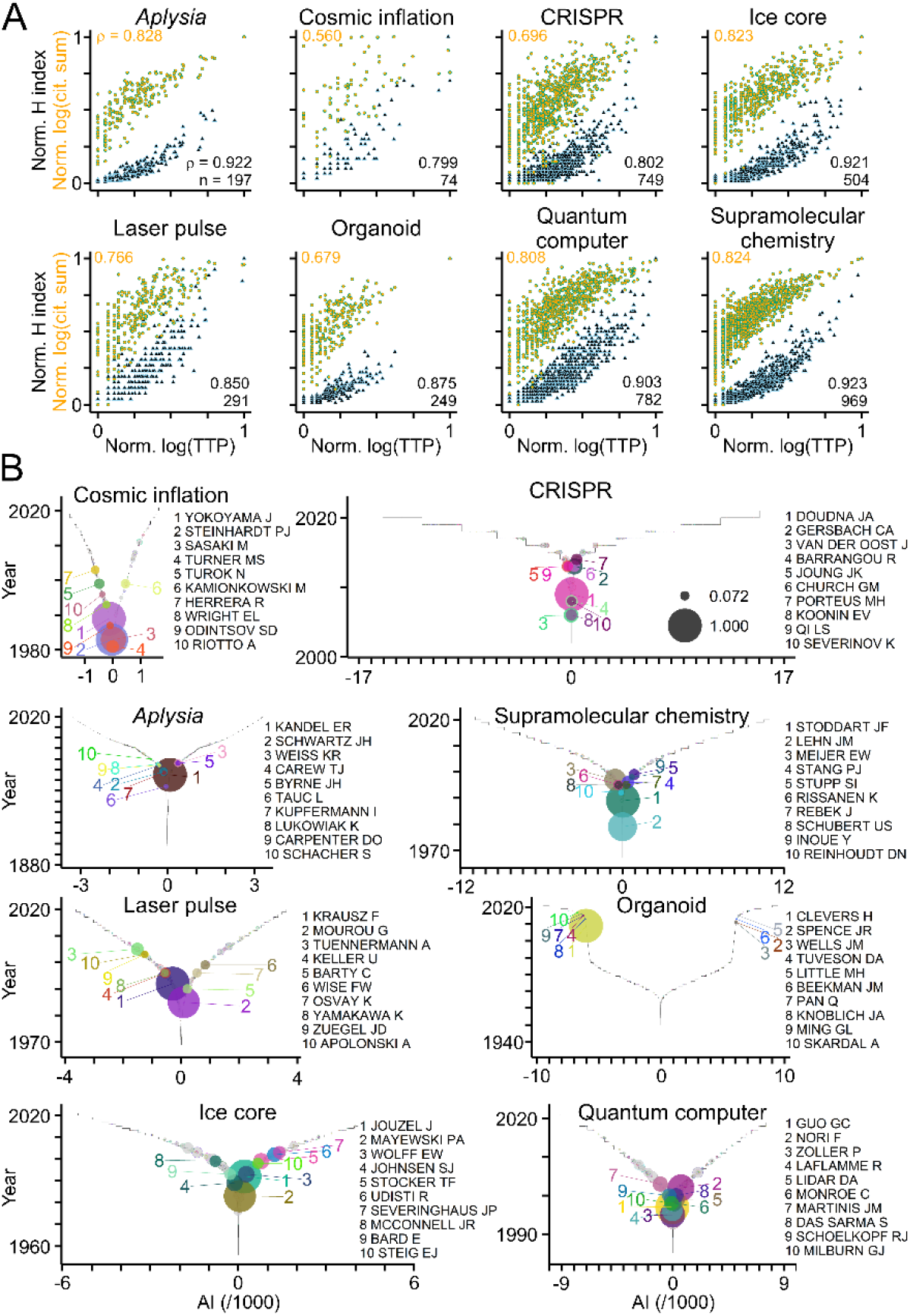
TTP-based evaluation across fields and disciplines. (A) Scatterplots where circles represent individual authors publishing in the selected fields of science and technology (Table 2) with their H indices (black-blue triangles; normalized to maximum) and sum of citations (orange-green circles; log10 values normalized to maximum) plotted against their TTP values (log10 values normalized to maximum). Numbers indicate rho values and sample sizes (Spearman’s correlation test; p < 0.0001 for all comparisons). (B) Graphs showing TTP values of authors in selected fields with colored circles and names indicating authors with ten highest TTP values. Grey circles with colored border indicate authors with TTP values above zero. Circle size indicates log10(TTP) normalized to maximum.

## Discussion

TTA fills a gap between global investigations of the scientific endeavour and the recurrent need to identify and evaluate the teams working on a user-defined topic of interest in science and technology.

A prime feature is the new measure to estimate scientific production named TTP. Several aspects distinguish this metric from existing author-level indicators. TTP takes into account three important aspects of research activity: the publication of peer-reviewed scientific articles, the training and mentoring of junior scientists, who continue their career within the field, and the establishment of collaborative connections that signify recognition due to specific expertise and capacities. The respective parameters are derived solely from the author(s) of scientific articles and the year of publication. Thus, TTP estimates scientific production independently from citation counts and augments the group of indicators that do not rely on this factor [46–49]. Notably, the significant correlation of TTP values of authors with their numbers of citations and their H indices in all fields tested indicates the usefulness of the new measure. A second feature introduced here are new visuals named TTGs that provide users with ad-hoc views on the workforce driving a field. They reveal its origin, development and size, and expose the publication records of authors as well as their genealogical and collaborative connections. These graphs complement present approaches to display bibliometric information and to visualize different aspects of scientific production [50–58].

TTA exposes factors that impact the workforce development of a field. For example, the calculation of publication periods revealed that few authors contributed for more than one year to the Clock field. This finding supports previous reports that in many research areas only a small fraction of the workforce publishes during long periods of time [59]. The delineation of families and collaborator networks in the Clock field revealed that genealogical and collaborative connections prolong the life-span of authors. These observations are in line with studies showing the relevance of training and mentorship [60–64] and the importance of collaborations [65–72]. The automatic delineation of family connections from first author-last author pairs provides an alternative to efforts requiring user input [73–75] (https://www.genealogy.math.ndsu.nodak.edu/, https://academictree.org/). However, TTA underestimates offspring counts in the case of co-first or co-last authorship, of alphabetical author lists or of field-specific author ranking [76, 77]. Other caveats should be mentioned: TTP values are field-specific, scale with the size of research groups and depend on the publication period of authors. Therefore, TTP-based ranking is context-dependent and unsuited to evaluate junior scientists [78]. Moreover, TTP is highly selective as only a fraction of authors has non-zero values, and it cannot value innovative, ground-breaking contributions from small teams or from teams that contribute only briefly to a field. TTA like all other name-dependent approaches faces the challenge of author disambiguation, which can be mitigated by assignment of unique author identifiers (https://orcid.org/) and computational algorithms [5, 29, 79–83]. Honorary and ghost authorship will confound results of TTA depending on their prevalence in the field [84, 85].

Peer-reviewed articles were used to introduce the features of TTA as this form of publication represents the core of scientific production [1], but the approach may also be applied to other types of publications such as preprints [86] and patents [87]. Future versions of TTA should provide web-based access to TTA allowing for direct retrieval and immediate processing of bibliographic information and the interactive display of results.

## Acknowledgments

The author thanks V. Demais, S. Eglen, N. Elghobashi-Meinhardt, J. Jouzel, J.M. Lehn, M. Muzet, V. Pallottini, J.L. Paluh, B. Poulain, H. Runz, J.P. Sauvage, D. Schulte, S. Silber and M. Slezak for helpful discussions and comments on previous versions of the manuscript.

## Supporting information

**S1 File. TTA-derived results for the Clock field.** Csv file summarizing TTA data for the Clock field using PubMed articles related to “circadian clock”.

## References

1. Tennant JP, Ross-Hellauer T. The limitations to our understanding of peer review. Research integrity and peer review. 2020;5:6. Epub 2020/05/06. doi: 10.1186/s41073-020-00092-1. PubMed PMID: 32368354; PubMed Central PMCID: PMCPMC7191707.

2. Bornmann L, Mutz R. Growth rates of modern science: A bibliometric analysis based on the number of publications and cited references. J Assoc Inf Sci Technol. 2015;66(11):2215–22. doi: 10.1002/asi.23329.

3. Clauset A, Larremore DB, Sinatra R. Data-driven predictions in the science of science. Science. 2017;355(6324):477–80. doi: 10.1126/science.aal4217. PubMed PMID: WOS:000393183100032.

4. Mukherjee S, Romero DM, Jones B, Uzzi B. The nearly universal link between the age of past knowledge and tomorrow’s breakthroughs in science and technology: The hotspot. Science advances. 2017;3(4):e1601315. Epub 2017/04/26. doi: 10.1126/sciadv.1601315. PubMed PMID: 28439537; PubMed Central PMCID: PMCPMC5397134.

5. Zeng A, Shen ZS, Zhou JL, Wu JS, Fan Y, Wang YG, et al. The science of science: From the perspective of complex systems. Phys Rep. 2017;714:1–73. doi: 10.1016/j.physrep.2017.10.001. PubMed PMID: WOS:000418464100001.

6. Fortunato S, Bergstrom CT, Borner K, Evans JA, Helbing D, Milojevic S, et al. Science of science. Science. 2018;359(6379). Epub 2018/03/03. doi: 10.1126/science.aao0185. PubMed PMID: 29496846; PubMed Central PMCID: PMCPMC5949209.

7. Fire M, Guestrin C. Over-optimization of academic publishing metrics: observing Goodhart’s Law in action. GigaScience. 2019;8(6). Epub 2019/05/31. doi: 10.1093/gigascience/giz053. PubMed PMID: 31144712; PubMed Central PMCID: PMCPMC6541803.

8. Hardwicke TE, Serghiou S, Janiaud P, Danchev V, Crüwell S, Goodman SN, et al. Calibrating the Scientific Ecosystem Through Meta-Research. Annu Rev Stat Appl. 2020;7(1):null. doi: 10.1146/annurev-statistics-031219-041104.

9. Harzing AW, Alakangas S. Google Scholar, Scopus and the Web of Science: a longitudinal and cross-disciplinary comparison. Scientometrics. 2016;106(2):787–804. doi: 10.1007/s11192-015-1798-9. PubMed PMID: WOS:0003 69017300015.

10. Gusenbauer M, Haddaway NR. Which academic search systems are suitable for systematic reviews or meta-analyses? Evaluating retrieval qualities of Google Scholar, PubMed, and 26 other resources. Research Synthesis Methods. 2020;11(2):181–217. doi: 10.1002/jrsm.1378. PubMed PMID: WOS:000509659900001.

11. Agarwal P, Searls DB. Can literature analysis identify innovation drivers in drug discovery? Nature Reviews Drug Discovery. 2009;8(11):865–78. doi: 10.1038/nrd2973. PubMed PMID: WOS:000271388200020.

12. Cunningham H, Tablan V, Roberts A, Bontcheva K. Getting More Out of Biomedical Documents with GATE’s Full Lifecycle Open Source Text Analytics. PLOS Computational Biology. 2013;9(2):e1002854. doi: 10.1371/journal.pcbi.1002854.

13. Chen Q, Lee K, Yan S, Kim S, Wei CH, Lu Z. BioConceptVec: Creating and evaluating literature-based biomedical concept embeddings on a large scale. PLoS Comput Biol. 2020;16(4):e1007617. Epub 2020/04/24. doi: 10.1371/journal.pcbi.1007617. PubMed PMID: 32324731; PubMed Central PMCID: PMCPMC7237030.

14. Venkatakrishnan AJ, Puranik A, Anand A, Zemmour D, Yao X, Wu X, et al. Knowledge synthesis of 100 million biomedical documents augments the deep expression profiling of coronavirus receptors. eLife. 2020;9:e58040. doi: 10.7554/eLife.58040.

15. Dridi A, Gaber MM, Azad RMA, Bhogal J. Scholarly data mining: A systematic review of its applications. Wiley Interdisciplinary Reviews-Data Mining and Knowledge Discovery. 2021;11(2). doi: 10.1002/widm.1395. PubMed PMID: WOS:0005 87964800001.

16. Rivest M, Vignola-Gagne E, Archambault E. Article-level classification of scientific publications: A comparison of deep learning, direct citation and bibliographic coupling. PloS one. 2021;16(5):e0251493–e. doi: 10.1371/journal.pone.0251493. PubMed PMID: MEDLINE:33974653.

17. Cronin B. Hyperauthorship: A postmodern perversion or evidence of a structural shift in scholarly communication practices? Journal of the American Society for Information Science and Technology. 2001;52(7):558–69. doi: 10.1002/asi.1097. PubMed PMID: WOS:000168426400005.

18. Claxton LD. Scientific authorship: Part 2. History, recurring issues, practices, and guidelines. Mutation Research/Reviews in Mutation Research. 2005;589(1):31–45. doi: https://doi.org/10.1016/j.mrrev.2004.07.002.

19. Marusic A, Bosnjak L, Jeroncic A. A Systematic Review of Research on the Meaning, Ethics and Practices of Authorship across Scholarly Disciplines. Plos One. 2011;6(9). doi: 10.1371/journal.pone.0023477. PubMed PMID: WOS:000294802800004.

20. Sauermann H, Haeussler C. Authorship and contribution disclosures. Science advances. 2017;3(11):e1700404. Epub 2017/11/21. doi: 10.1126/sciadv.1700404. PubMed PMID: 29152564; PubMed Central PMCID: PMCPMC5687853.

21. Holcombe AO. Contributorship, Not Authorship: Use CRediT to Indicate Who Did What. Publications. 2019;7(3). doi: 10.3390/publications7030048. PubMed PMID: WOS:000487987700001.

22. Hicks D, Wouters P, Waltman L, de Rijcke S, Rafols I. The Leiden Manifesto for research metrics. Nature. 2015;520(7548):429–31. doi: 10.1038/520429a. PubMed PMID: WOS:000353334500013.

23. Hirsch JE. An index to quantify an individual’s scientific research output. Proceedings of the National Academy of Sciences of the United States of America. 2005;102(46):16569–72. Epub 2005/11/09. doi: 10.1073/pnas.0507655102. PubMed PMID: 16275915; PubMed Central PMCID: PMCPMC1283832.

24. Garfield E. The History and Meaning of the Journal Impact Factor. JAMA. 2006;295(1):90–3. doi: 10.1001/jama.295.1.90.

25. Docampo D, Bessoule J-J. A new approach to the analysis and evaluation of the research output of countries and institutions. Scientometrics. 2019;119(2):1207–25. doi: 10.1007/s11192-019-03089-w.

26. Waltman L. A review of the literature on citation impact indicators. Journal of Informetrics. 2016;10(2):365–91. doi: 10.1016/j.joi.2016.02.007. PubMed PMID: WOS:000377413800004.

27. Aksnes DW, Langfeldt L, Wouters P. Citations, Citation Indicators, and Research Quality: An Overview of Basic Concepts and Theories. Sage Open. 2019;9(1). doi: 10.1177/2158244019829575. PubMed PMID: WOS:000458649200001.

28. Braithwaite J, Herkes J, Churruca K, Long JC, Pomare C, Boyling C, et al. Comprehensive Researcher Achievement Model (CRAM): a framework for measuring researcher achievement, impact and influence derived from a systematic literature review of metrics and models. BMJ Open. 2019;9(3):e025320. doi: 10.1136/bmjopen-2018-025320.

29. Milojevic S. Accuracy of simple, initials-based methods for author name disambiguation. Journal of Informetrics. 2013;7(4):767–73. doi: 10.1016/j.joi.2013.06.006. PubMed PMID: WOS:000327920400001.

30. Newman ME. The structure of scientific collaboration networks. Proceedings of the National Academy of Sciences of the United States of America. 2001;98(2):404–9. Epub 2001/01/10. doi: 10.1073/pnas.021544898. PubMed PMID: 11149952; PubMed Central PMCID: PMCPMC14598.

31. R Core Team. R: A Language and Environment for Statistical Computing. Vienna, Austria: R Foundation for Statistical Computing; 2019.

32. Dowle M. data.table: Extension of ‘data.frame’ 2019. Available from: https://CRAN.R-project.org/package=data.table.

33. Csardi GN, T. The igraph software package for complex network research. InterJournal. 2006;Complex Systems:1695.

34. Dinno A. dunn.test: Dunn’s Test of Multiple Comparisons Using Rank Sums. 2017.

35. Larsson J. eulerr: Area-Proportional Euler and Venn Diagrams with Ellipses. 2019.

36. Tang YH, M.; Li, W. ggfortify: Unified Interface to Visualize Statistical Result of Popular R Packages. The R Journal. 2016;8(2).

37. Wickham H. Ggplot2: Elegant Graphics for Data Analysis. New York: Springer-Verlag; 2016.

38. Slowikowski KS, A.; Hughes, S.; Lukauskas, S.; Irisson, J.O.; Kamvar, Z.N.; Thompson, R.; Dervieux, C.; Yutani, H.; Gramme, P. ggrepel: Automatically Position Non-Overlapping Text Labels with ‘ggplot2’. 2018.

39. Soetaert K. plot3D: Plotting Multi-Dimensional Data. 2017.

40. Moroz LL. Aplysia. Current biology: CB. 2011;21(2):R60–1. Epub 2011/01/25. doi: 10.1016/j.cub.2010.11.028. PubMed PMID: 21256433; PubMed Central PMCID: PMCPMC4024469.

41. Hille F, Richter H, Wong SP, Bratovič M, Ressel S, Charpentier E. The Biology of CRISPR-Cas: Backward and Forward. Cell. 2018;172(6):1239–59. doi: https://doi.org/10.1016/j.cell.2017.11.032.

42. Simian M, Bissell MJ. Organoids: A historical perspective of thinking in three dimensions. The Journal of cell biology. 2017;216(1):31–40. Epub 12/28. doi: 10.1083/jcb.201610056. PubMed PMID: 28031422.

43. Lancaster MA, Knoblich JA. Organogenesis in a dish: Modeling development and disease using organoid technologies. Science. 2014;345(6194):1247125. doi: 10.1126/science.1247125.

44. Garreta E, Kamm RD, Chuva de Sousa Lopes SM, Lancaster MA, Weiss R, Trepat X, et al. Rethinking organoid technology through bioengineering. Nature Materials. 2021;20(2):145–55. doi: 10.1038/s41563-020-00804-4.

45. Patke A, Young MW, Axelrod S. Molecular mechanisms and physiological importance of circadian rhythms. Nature Reviews Molecular Cell Biology. 2020;21(2):67–84. doi: 10.1038/s41580-019-0179-2.

46. Liu X, Bollen J, Nelson ML, Van De Sompel H, Egghe L. Co-authorship networks in the digital library research community. Infometrics. 2005;41(6):1462–80. PubMed PMID: edsfra.17035814.

47. Sugimoto CR, Work S, Lariviere V, Haustein S. Scholarly Use of Social Media and Altmetrics: A Review of the Literature. Journal of the Association for Information Science and Technology. 2017;68(9):2037–62. doi: 10.1002/asi.23833. PubMed PMID: WOS:000407793000001.

48. Bornmann L, Haunschild R. Normalization of zero-inflated data: An empirical analysis of a new indicator family and its use with altmetrics data. Journal of Informetrics. 2018;12(3):998–1011. doi: 10.1016/j.joi.2018.01.010. PubMed PMID: WOS:000442670600029.

49. Tahamtan I, Bornmann L. Altmetrics and societal impact measurements: Match or mismatch? A literature review. Profesional De La Informacion. 2020;29(1). doi: 10.3145/epi.2020.ene.02. PubMed PMID: WOS:000531809400002.

50. Börner K, Chen C, Boyack KW. Visualizing knowledge domains. Annual Review of Information Science & Technology. 2003;37(1):179–255. doi: 10.1002/aris.1440370106. PubMed PMID: 64982071.

51. Chen C. Citespace II: Detecting and visualizing emerging trends and transient patterns in scientific literature. Journal of the American Society for Information Science and Technology (Print). 2006;57(3):359–77. PubMed PMID: edsfra.17473696.

52. van Eck NJ, Waltman L. Software survey: VOSviewer, a computer program for bibliometric mapping. Scientometrics. 2010;84(2):523–38. doi: 10.1007/s11192-009-0146-3.

53. Cobo MJ, Lopez-Herrera AG, Herrera-Viedma E, Herrera F. SciMAT: A new science mapping analysis software tool. Journal of the American Society for Information Science and Technology. 2012;63(8):1609–30. doi: 10.1002/asi.22688. PubMed PMID: WOS:000306758600010.

54. Marx W, Bornmann L, Barth A, Leydesdorff L. Detecting the Historical Roots of Research Fields by Reference Publication Year Spectroscopy (RPYS). Journal of the Association for Information Science & Technology. 2014;65(4):751–64. doi: 10.1002/asi.23089. PubMed PMID: 94969748.

55. van Eck NJ, Waltman L. CitNetExplorer: A new software tool for analyzing and visualizing citation networks. 2014. p. 802.

56. Aria M, Cuccurullo C. bibliometrix: An R-tool for comprehensive science mapping analysis. Journal of Informetrics. 2017;11(4):959–75. doi: https://doi.org/10.1016/j.joi.2017.08.007.

57. Bornmann L, Haunschild R. Plots for visualizing paper impact and journal impact of single researchers in a single graph. Scientometrics. 2018;115(1):385–94. doi: 10.1007/s11192-018-2658-1. PubMed PMID: WOS:000426807700020.

58. Moral-Munoz JA, Herrera-Viedma E, Santisteban-Espejo A, Cobo MJ. Software tools for conducting bibliometric analysis in science: An up-to-date review. Profesional De La Informacion. 2020;29(1). doi: 10.3145/epi.2020.ene.03. PubMed PMID: WOS:000531809400004.

59. Ioannidis JP, Boyack KW, Klavans R. Estimates of the continuously publishing core in the scientific workforce. PLoS One. 2014;9(7):e101698. Epub 2014/07/10. doi: 10.1371/journal.pone.0101698. PubMed PMID: 25007173; PubMed Central PMCID: PMCPMC4090124.

60. Malmgren RD, Ottino JM, Nunes Amaral LA. The role of mentorship in protégé performance. Nature. 2010;465(7298):622–6. doi: 10.1038/nature09040.

61. Lienard JF, Achakulvisut T, Acuna DE, David SV. Intellectual synthesis in mentorship determines success in academic careers. Nature communications. 2018;9(1):4840. Epub 2018/11/30. doi: 10.1038/s41467-018-07034-y. PubMed PMID: 30482900; PubMed Central PMCID: PMCPMC6258699.

62. Sekara V, Deville P, Ahnert SE, Barabasi AL, Sinatra R, Lehmann S. The chaperone effect in scientific publishing. Proceedings of the National Academy of Sciences of the United States of America. 2018;115(50):12603–7. Epub 2018/12/12. doi: 10.1073/pnas.1800471115. PubMed PMID: 30530676; PubMed Central PMCID: PMCPMC6294962.

63. Li W, Aste T, Caccioli F, Livan G. Early coauthorship with top scientists predicts success in academic careers. Nature communications. 2019;10(1):5170. Epub 2019/11/16. doi: 10.1038/s41467-019-13130-4. PubMed PMID: 31729362; PubMed Central PMCID: PMCPMC6858367.

64. Ma YF, Mukherjee S, Uzzi B. Mentorship and protege success in STEM fields. Proceedings of the National Academy of Sciences of the United States of America. 2020;117(25):14077–83. doi: 10.1073/pnas.1915516117. PubMed PMID: WOS:000546772500009.

65. Melin G, Persson O. Studying research collaboration using co-authorships. Scientometrics. 1996;36(3):363–77. doi: 10.1007/bf02129600. PubMed PMID: WOS:A1996VG50500006.

66. Barabási AL, Jeong H, Néda Z, Ravasz E, Schubert A, Vicsek T. Evolution of the social network of scientific collaborations. Physica A: Statistical Mechanics and its Applications. 2002;311(3):590–614. doi: https://doi.org/10.1016/S0378-4371(02)00736-7.

67. Wuchty S, Jones BF, Uzzi B. The increasing dominance of teams in production of knowledge. Science. 2007;316(5827):1036–9. Epub 2007/04/14. doi: 10.1126/science.1136099. PubMed PMID: 17431139.

68. Milojevic S. Principles of scientific research team formation and evolution. Proceedings of the National Academy of Sciences of the United States of America. 2014;111(11):3984–9. Epub 2014/03/05. doi: 10.1073/pnas.1309723111. PubMed PMID: 24591626; PubMed Central PMCID: PMCPMC3964124.

69. Lariviere V, Gingras Y, Sugimoto CR, Tsou A. Team size matters: Collaboration and scientific impact since 1900. Journal of the Association for Information Science and Technology. 2015;66(7):1323–32. doi: 10.1002/asi.23266. PubMed PMID: WOS:000355858200002.

70. Fonseca Bde P, Sampaio RB, Fonseca MV, Zicker F. Co-authorship network analysis in health research: method and potential use. Health research policy and systems. 2016;14(1):34. Epub 2016/05/04. doi: 10.1186/s12961-016-0104-5. PubMed PMID: 27138279; PubMed Central PMCID: PMCPMC4852432.

71. Parish AJ, Boyack KW, Ioannidis JPA. Dynamics of co-authorship and productivity across different fields of scientific research. PLoS One. 2018;13(1):e0189742. Epub 2018/01/11. doi: 10.1371/journal.pone.0189742. PubMed PMID: 29320509; PubMed Central PMCID: PMCPMC5761855.

72. Ahmadpoor M, Jones BF. Decoding team and individual impact in science and invention. Proceedings of the National Academy of Sciences of the United States of America. 2019;116(28):13885–90. Epub 2019/06/27. doi: 10.1073/pnas.1812341116. PubMed PMID: 31235568; PubMed Central PMCID: PMCPMC6628781.

73. David SV, Hayden BY. Neurotree: a collaborative, graphical database of the academic genealogy of neuroscience. PLoS One. 2012;7(10):e46608. Epub 2012/10/17. doi: 10.1371/journal.pone.0046608. PubMed PMID: 23071595; PubMed Central PMCID: PMCPMC3465338.

74. Hirshman BR, Tang JA, Jones LA, Proudfoot JA, Carley KM, Marshall L, et al. Impact of medical academic genealogy on publication patterns: An analysis of the literature for surgical resection in brain tumor patients. Annals of neurology. 2016;79(2):169–77. Epub 2016/01/05. doi: 10.1002/ana.24569. PubMed PMID: 26727354.

75. Sanyal DK, Dey S, Das PP. g(m)-index: a new mentorship index for researchers. Scientometrics. 2020;123(1):71–102. doi: 10.1007/s11192-020-03384-x. PubMed PMID: WOS:000516437800003.

76. Frandsen TF, Nicolaisen J. What is in a name? Credit assignment practices in different disciplines. Journal of Informetrics. 2010;4(4):608–17. doi: 10.1016/j.joi.2010.06.010. PubMed PMID: WOS:000281616200015.

77. Waltman L. An empirical analysis of the use of alphabetical authorship in scientific publishing. Journal of Informetrics. 2012;6(4):700–11. doi: 10.1016/j.joi.2012.07.008. PubMed PMID: WOS:000308581700029.

78. Schimanski LA, Alperin JP. The evaluation of scholarship in academic promotion and tenure processes: Past, present, and future. F1000Research. 2018;7:1605-. doi: 10.12688/f1000research.16493.1. PubMed PMID: MEDLINE:30647909.

79. Torvik VI, Smalheiser NR. Author Name Disambiguation in MEDLINE. Acm Transactions on Knowledge Discovery from Data. 2009;3(3). doi: 10.1145/1552303.1552304. PubMed PMID: WOS:000208168000001.

80. Tang L, Walsh JP. Bibliometric fingerprints: name disambiguation based on approximate structure equivalence of cognitive maps. Scientometrics. 2010;84(3):763–84. doi: 10.1007/s11192-010-0196-6. PubMed PMID: WOS:000280274400014.

81. D’Angelo CA, Giuffrida C, Abramo G. A Heuristic Approach to Author Name Disambiguation in Bibliometrics Databases for Large-Scale Research Assessments. Journal of the American Society for Information Science and Technology. 2011;62(2):257–69. doi: 10.1002/asi.21460. PubMed PMID: WOS:000286687300005.

82. Glänzel W, Heeffer S, Thijs B. A triangular model for publication and citation statistics of individual authors. Scientometrics. 2016;107(2):857–72. doi: 10.1007/s11192-016-1870-0.

83. Albert PJ, Dutta S, Lin J, Zhu ZM, Bales M, Johnson SB, et al. ReCiter: An open source, identity-driven, authorship prediction algorithm optimized for academic institutions. Plos One. 2021;16(4). doi: 10.1371/journal.pone.0244641. PubMed PMID: WOS:000636467000022.

84. Wislar JS, Flanagin A, Fontanarosa PB, Deangelis CD. Honorary and ghost authorship in high impact biomedical journals: a cross sectional survey. BMJ (Clinical research ed). 2011;343:d6128. Epub 2011/10/27. doi: 10.1136/bmj.d6128. PubMed PMID: 22028479; PubMed Central PMCID: PMCPMC3202014.

85. Al-Herz W, Haider H, Al-Bahhar M, Sadeq A. Honorary authorship in biomedical journals: how common is it and why does it exist? Journal of medical ethics. 2014;40(5):346–8. Epub 2013/08/21. doi: 10.1136/medethics-2012-101311. PubMed PMID: 23955369.

86. Abdill RJ, Blekhman R. Tracking the popularity and outcomes of all bioRxiv preprints. eLife. 2019;8. doi: 10.7554/eLife.45133. PubMed PMID: WOS:000467693500001.

87. Sick N, Merig JM, Kratzig O, List J. Forty years of World Patent Information: A bibliometric overview. World Patent Information. 2021;64. doi: 10.1016/j.wpi.2020.102011. PubMed PMID: WOS:000636276000004.

